# Codon usage pattern reveals SARS-CoV-2 as a monomorphic pathogen of hybrid origin with role of silent mutations in rapid evolutionary success

**DOI:** 10.1101/2020.10.12.335521

**Authors:** Kanika Bansal, Sanjeet Kumar, Prabhu B. Patil

**Affiliations:** CSIR- Institute of Microbial Technology, Chandigarh, India; Gangadhar Meher University, Odisha, India

## Abstract

Viruses are dependent on the host tRNA pool, and an optimum codon usage pattern (CUP) is the driving force in its evolution. Systematic analysis of CUP of the coding sequences (CDS) of representative major pangolin lineages A and B of SARS-CoV-2 indicate a single transmission event of a codon-optimized virus from its source into humans. Here, no direct congruence could be detected in CUP of all CDS of SARS-CoV-2 with the non-human natural SARS viruses further reiterating its novelty. Several CDS show similar CUP with bat or pangolin, while others have distinct CUP pointing towards a possible hybrid nature of the virus. At the same time, phylogenetic diversity suggests the role of even silent mutations in its success by adapting to host tRNA pool. However, genomes of SARS-CoV-2 from primary infections are required to investigate the origins amongst the competing natural or lab leak theories.

## Introduction

The origin and success of the novel SARS coronavirus (SARS-CoV-2) (betacoronavirus) causing the COVID-19 disease pandemic has been a topic of intense discussion. In the past two decades since the first outbreak of SARS in 2002, several SARS-related coronaviruses were reported from the bat, which was speculated to be significant reservoir for future possible outbreaks ^1–4^. Bats are the only flying mammals representing 20% of known mammalian species and are critical natural reservoirs of many zoonotic viruses like Nipah virus, Hendra virus, rabies virus, Ebola virus, etc. ^5–7^. Besides bat, a considerable number of wild animals have played a pivotal role in zoonotic transfers ^8^. According to reports before SARS-CoV-2 pandemic, due to human interventions, there was a high-risk assessment of SARS coronavirus infection from wild animals like bats, civets, pangolins, snakes, tiger and primates in China ^9–11^.

The human-wildlife interface as a part of culture or globalization poses risks for zoonotic transfers followed by disease outbreaks like coronavirus outbreaks: SARS (2002,2003), MERS (2012), and SARS-CoV-2 (2019). Animal reservoirs for such outbreaks are estimated by their genome similarities with already reported SARS viruses from diverse animals. For instance, the SARS 2003 outbreak virus had 99.6% genome similarity with palm civets indicating it to be a direct source. Just 0.4% divergence from the animal reservoir stipulates its recent transfer into the masked palm civet population ^12^. Despite genetic diversity with bat SARS-CoV they were ultimately found to be a source of the pandemic due to no pathogen prevalence in wild civet population and clinical symptom manifestation in civets, unlike bats ^4^. In the current pandemic, there are several theories of the origin of SARS-CoV-2 either from bat, pangolin, dog, or some intermediate host, etc. ^13, 14^. The closest match to SARS-CoV-2 is RaTG13 (96% identity), isolated from *Rhinolophus affinis* bat ^15^, followed by pangolin SARS viruses with 91% identity ^16^. As the closest match is just 96%, it has opened a heated debate in the scientific community for its origin, and no direct animal source can be detected.

According to genome similarities, SARS-CoV-2 differs from its closest SARS coronavirus by 4%, followed by 9% with its next closest relative pangolin. It indicates that the virus has evolved before infecting humans, and there is a missing link between bat/pangolin and humans, which further inflates the argument on the animal source. Nevertheless, another study based on CpG island deficiency in SARS-CoV-2 and canine coronavirus (alphacoronavirus) suggested that dogs may have provided a cellular environment for SARS-CoV-2 evolution into a CpG deficient virus ^17^. Hence, they claim dog to be a direct source of the current pandemic, raising a constant debate ^18^ (https://www.linkedin.com/pulse/where-dog-laymans-version-my-mbe-paper-xuhua-xia/). But most other RNA viruses like pestvirus in addition to bat or pangolin SARS-CoV are also depleted in CpG are not included in the study. CpG island deficiency is not a unique feature of dog SARS-CoV and a later study contradicted that there is no direct evidence for the role of dogs as intermediate hosts ^18^.

Usage patterns of synonymous codons are a critical feature in the adaptation of organisms as viruses are dependent on the host tRNA pool for replication and disease manifestations. For instance, codon adaptation indices were studied for retroviruses infecting humans, including the HIV-1 virus ^19^. Once the viral genome is in the host translational mechanism, genes having optimized codons according to the host translate faster, resulting in higher fitness of the virus ^20^. Hence, for host jump events, viral codon optimization based on the host tRNA pool is critical ^21–23^. In the present study, we have focused on the codon usage pattern (CUP) of CDS of SARS coronavirus from different hosts under debate (bat, pangolin, and dog) as a probable origin for SARS-CoV-2. An optimum CUP is vital in its evolution, and probable host jumps, and this also results in synonymous changes in the viral genome, which are not revealed by mutational studies at protein level. Population based mutational analysis of SASR-CoV-2 at nucleotide level have revealed various silent mutations conserved in the genome ^24, 25^. These silent mutations may have consequent alteration in codon usage or translation efficiency (Mercatelli and Giorgi 2020). Systematic insight into CUP is required to trace the evolutionary trajectory, understand its origin, and remarkable success of emergent viruses like SARS-CoV-2. For a virus to be successful, it should be able to efficiently transmit to the host, i.e., recognize host (SARS-CoV-2 spike recognizing human ACE2), and once inside the host, it should replicate (RNA dependent RNA polymerase rdrp (ORF 1ab)) its ORFs (ORF 1ab, spike (S), ORF 3a, envelope (E), membrane (M), ORF 6, ORF 7a, ORF 7b, ORF 8 and nucleocapsid (N)). The rdrp and spike are now considered important targets for vaccine development for SARS-CoV-2 ^26–28^. Evolutionary studies till now suggest that the spike receptor-binding domain of SARS-CoV-2 is more similar to pangolin SARS strains as compared to bat SARS ^18^.

Presently, we have analyzed CUP of coding regions in SARS coronavirus isolates reported from humans, bat, pangolin, and dog. Here, we have calculated the percentage of GC biased synonymous codons for amino acids having at least four synonymous codons (Glycine, Valine, Threonine, Leucine, Arginine, Serine, Proline and Alanine) ^29^. Patil et. al, have proven how CUP can detect horizontally acquired genes from a diverse background under selection pressure by analyzing codon usage pattern of each amino acid in a particular gene in a graphical way. Similarly, a host jump event may lead to codon optimization, which will be reflected in the CUP. Ideally, for an organism, its crucial genes should have a similar pattern of CUP. Nevertheless, genes pivotal for viral host jump and disease manifestations like spike or rdrp may show deviation from the pattern. Hence, CUP graphs enable us to visually inspect the patterns of synonymous changes across diverse hosts and be suitable for addressing the surprising origin of the virus.

Interestingly, CUP for all the CDS for 134 SARS-CoV-2 genomes (supplementary table 1) was not diversified irrespective of their diverse phylogenetic lineages known in the population. This indicates recent and one-time introduction of an isolate into the human host. While diversity in phylogeny as seen by major and minor lineages suggests that even silent or synonymous mutations play an important role in the rapid emergence and spread. In this context, it is pertinent to note that any mutation can have a consequence in virus. It is dependent on tRNA pool of host that are biased towards a particular set of degenerate codons for a particular amino acid. In fact, a silent mutation can be lethal for a virus if matching tRNA is not encoded in the genome of host or absent in a particular cell or tissue. Further, studies in this regard need of the hour to understand this silent co-evolution in viruses in general and SARS-CoV-2 in particular. On the other hand, the CUPs for the CDSs for other probable hosts i.e. bat, dog and pangolins were diversified (as depicted from standard deviation bars in figure 1). Unlike SARS-CoV-2 isolates of humans, CUP of all CDS were variable in isolates of non-human hosts like a bat, pangolin, and dog (supplementary table 2) depicting ongoing adaptation and evolution of SARS in these hosts (figure 1). Amongst the non-human hosts, bat has most varied CUP correlating with the well-known fact that bat is a reservoir of SARS coronaviruses.

**Figure 1:**
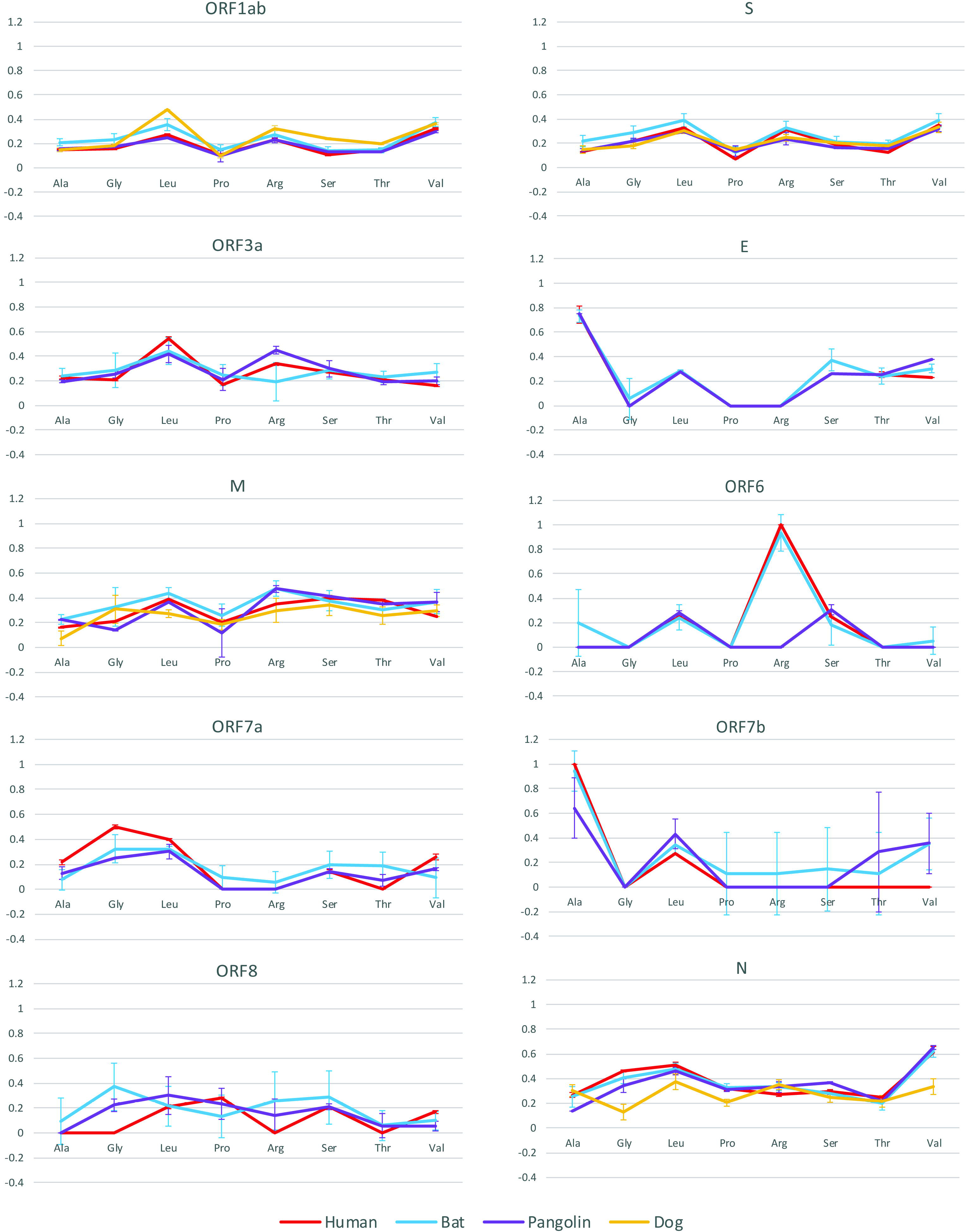
Codon usage pattern for all CDS of SARS genomes from human (SARS-CoV-2), bat, pangolin and dog. Eight amino acids with at least four synonymous codons are represented in the X-axis, and the percentage of codons ending with G/C for each amino acid is represented on Y-axis. Standard deviations for each amino acid codon usage is represented by vertical error bars.

Overall, CUP of SARS-CoV-2 for ORF 1ab, envelope and ORF 6, were overlapping with that of SARS from non-human hosts i.e., bat and pangolin, with some exceptions. For instance, ORF 1ab has overlapping pattern for SARS of pangolin origin with human SARS-CoV-2. Here, bat SARS also had similar pattern with SARS-CoV-2 with slightly higher fractions of codon usage for leucine and proline. Envelope protein had overlapping patterns of CUP of SARS-CoV-2 with SARS from pangolin and bat except for a slightly higher fraction of serine (bat SARS) and valine (bat and pangolin SARS). In case of ORF 6 also CUP of bat and pangolin have overlapping patterns with SARS-CoV-2. Here, pangolin SARS ORF 6 did not have arginine codons ending with G or C while, bat and SARS-CoV-2 had the maximum fraction (i.e. 1) of these. Further, CUP of SARS-CoV-2 for spike, ORF 7a, ORF 7b and nucleocapsid proteins were having similar pattern with that of SARS from bat or pangolin. However, CUP for ORF 3a, membrane and ORF 8 had distinct CUP patterns for SARS-CoV-2 compared with that of bat and pangolin. However, CUP of SARS from dog had distinct patterns for the CDS analysed in the study, clearly overruling dog as a probable source compared to bat and pangolin.

CUP of all CDS among lineages A and B were not diversified, indicating a single event of transmission of a codon-optimized SARS strain to the human population from its source. Further, CUP of SARS-CoV-2 is not showing congruency with its non-human natural counterparts. Hence, in the current study, we could not find closest relative of SARS-CoV-2 in natural settings which is in accordance to the previous genome similarity assessment. CUP pattern of ORF 1ab, envelope protein and ORF 6 is overlapping and spike protein, ORF 7a, ORF 7b and nucleocapsid protein is showing similar pattern, while, CUP of membrane protein, ORF 3a and ORF 8 are distinct from SARS of non-human hosts (bat or pangolin). It indicates that the evolution of all CDS is not linked. It can be depicted that either SARS-CoV-2 is a hybrid virus or the closest relative in natural settings is not yet discovered.

However, lack of closely related natural source of SARS-CoV-2 have now shifted lab leak theory to the mainstream from the conspiracy theory ^30, 31^ Hence, the probable origin of SARS-CoV-2 is a debate between two competing hypotheses of natural or lab leak. In order to find the true origin, we need to include SARS-CoV-2 from primary infection cases.

## Supplementary material

**Supplementary table 1:** Metadata for the human SARS-CoV-2 genomes used in the study

**Supplementary table 2:** SARS strains from non-human hosts used in the present study.

